# Patchouli alcohol suppresses gastric cancer growth and immune evasion via inhibition of the NF-κB/PD-L1 axis

**DOI:** 10.64898/2026.03.17.712304

**Authors:** Kun Hou, Qin Hao, Hao Yang, Fengxue Dai, Xuying Wang, Yu wei Dai, Li Feng, Haiwen Lu, Zhenfei Wang

**Affiliations:** Department of Pharmacy, Peking University Cancer Hospital (Inner Mongolia Campus)/Affiliated Cancer Hospital of Inner Mongolia Medical University, Hohhot, Inner Mongolia, China; The Central Laboratory for Digestive Tumor, Peking University Cancer Hospital (Inner Mongolia Campus)/Affiliated Cancer Hospital of Inner Mongolia Medical University, Hohhot, Inner Mongolia, China; The Laboratory for Inheritance and Development of Integrated Chinese (Mongolian) and Western Medicine in Anti-Tumor Therapy, Peking University Cancer Hospital (Inner Mongolia Campus)/Affiliated Cancer Hospital of Inner Mongolia Medical University, Hohhot, Inner Mongolia, China; Department of Gastrointestinal Surgery, Affiliated Hospital of Inner Mongolia Medical University, Hohhot, Inner Mongolia, China; Department of Radiation Oncology, Peking University Cancer Hospital (Inner Mongolia Campus)/Affiliated Cancer Hospital of Inner Mongolia Medical University, Hohhot, Inner Mongolia, China; Department of Day Chemotherapy Ward, Peking University Cancer Hospital (Inner Mongolia Campus)/Affiliated Cancer Hospital of Inner Mongolia Medical University, Hohhot, Inner Mongolia, China; Graduate School, Inner Mongolia Medical University, Hohhot, Inner Mongolia, China; Department of Medical Simulated Center, Inner Mongolia Medical University, Hohhot, Inner Mongolia, China

**Keywords:** Patchouli alcohol (PA), gastric cancer, immune evasion, NF-κB/PD-L1

## Abstract

**Objective:** This study aimed to investigate the anti-gastric cancer effect of Patchouli alcohol (PA), especially its influence on PD-L1-mediated immune evasion, and to elucidate the underlying molecular mechanisms.

**Methods:** A CCK-8 assay was used to evaluate the effects of PA on the viability of the gastric cancer cell lines HGC-27 and MKN-45. RT‒qPCR and western blotting were performed to analyze the mRNA and protein levels of NF-κB and PD-L1, respectively. In a coculture system of gastric cancer cells and peripheral blood mononuclear cells (PBMCs), the effect of PA pretreatment on the PBMC-induced apoptosis of cancer cells was analyzed by flow cytometry, and the cytotoxic activity of the PBMCs was assessed by a lactate dehydrogenase (LDH) release assay. Flow cytometry was also used to determine the proportions of CD3⁺CD8⁺ T cells and IFN-γ⁺CD8⁺ T cells. ELISA was used to measure the levels of IFN-γ, TNF-α, and granzyme B in the coculture supernatants. Immunofluorescence staining was conducted to assess NF-κB nuclear translocation. In a mouse xenograft model, tumor volume and weight were measured after 14 days of PA treatment. Histopathological changes and apoptosis were analyzed by HE and TUNEL staining. A luciferase reporter assay was used to examine the transcriptional regulation of PD-L1 by NF-κB.

**Results:** PA inhibited the viability of HGC-27 and MKN-45 cells in a dose- and time-dependent manner and significantly downregulated the expression of NF-κB and PD-L1 at both the mRNA and protein levels. In a PBMC coculture model, PA pretreatment enhanced the ability of PBMCs to induce apoptosis and directly kill gastric cancer cells. Furthermore, PA pretreatment increased the proportions of CD3⁺CD8⁺ T cells and IFN-γ⁺CD8⁺ T cells in a dose-dependent manner. Consistent with this immunostimulatory effect, PA increased the levels of IFN-γ, TNF-α, and granzyme B in the coculture supernatants. Mechanistically, western blotting analysis demonstrated that PA significantly reduced the protein levels of AKT, NF-κB, and PD-L1 in gastric cancer cells. Immunofluorescence staining further indicated that PA suppressed the nuclear translocation of NF-κB. In a mouse xenograft model, PA treatment significantly inhibited tumor growth, induced apoptosis, and downregulated NF-κB and PD-L1 protein expression in tumor tissues. Flow cytometry of tumor-infiltrating lymphocytes revealed increased proportions of CD3⁺CD8⁺ and IFN-γ⁺CD8⁺ T cells following PA treatment. Finally, luciferase reporter assays demonstrated that NF-κB directly regulates PD-L1 transcription by binding to its promoter region.

**Conclusion:** PA exerts antitumor effects in gastric cancer by suppressing the NF-κB/PD-L1 axis, thereby enhancing CD8⁺ T-cell–mediated cytotoxicity and inhibiting immune evasion.

## 1 INTRODUCTION

Gastric cancer (GC) ranks as the fifth most common malignancy and the third leading cause of cancer-related death globally (1). It was projected that by 2024, nearly one million new cases would be diagnosed annually worldwide, with China bearing approximately 40% of this burden. The established risk factors for gastric cancer include *Helicobacter pylori* infection, high dietary salt intake, and genetic susceptibility (2). Given the limited efficacy of existing therapies, the development of novel and more effective treatment strategies is urgently needed.

In recent years, immune checkpoint inhibitors (ICIs) have demonstrated remarkable efficacy across multiple solid tumors, including GC. Landmark global phase III trials, such as CheckMate–649 (3), Attraction–4 (4), KEYNOTE–859 (5)(5), ORIENT–16 (6), and RATIONALE–305 (7, 8), have led to the approval of PD–1 inhibitors (e.g., pembrolizumab and nivolumab) and PD–L1 inhibitors (e.g., sugemalimab) for the treatment of advanced or metastatic GC. Moreover, PD-1/PD-L1 inhibitors combined with chemotherapy have become the new standard first-line treatment, substantially improving clinical outcomes (9). These findings underscore that blockade of the PD-1/PD-L1 axis is particularly effective in PD-L1–positive GC patients, with higher PD-L1 expression levels correlated with greater therapeutic benefit. However, the broader application of ICIs faces challenges, including high cost, restricted insurance coverage, and limited accessibility, as well as immune-related adverse events (irAEs) that may lead to treatment interruption or premature discontinuation.

In contrast to ICIs, traditional Chinese medicine (TCM) has attracted increasing attention because of its favorable safety profile and multitarget synergistic potential. Accumulating *in vitro* and *in vivo* studies indicate that various TCM-derived monomers and formulae can modulate the PD–1/PD–L1 axis, remodel the tumor immune microenvironment, and enhance effector T-cell function, thereby not only potentiating immunotherapy but also mitigating certain irAEs (10). Notable examples include ginsenosides (11), astragalus polysaccharides (12), and berberine (13), which downregulate PD–L1 expression or suppress its downstream signaling. Moreover, several classic formulae have been demonstrated to delay immune exhaustion and improve antitumor immune responses (14,15).

Patchouli (Pogostemon cablin) is a traditional medicinal herb known for its spleen-strengthening, anti-inflammatory, and immunomodulatory properties. Its principal bioactive component, patchouli alcohol (PA), displays broad antitumor activity and suppresses proliferation and invasion in multiple cancer types. For instance, PA inhibits castration-resistant prostate cancer (CRPC) progression by blocking the NF-κB/Mcl-1 axis, inducing mitochondrial dysfunction, and downregulating the expression of metastasis-related markers (MMPs/VEGFs) (16). In non-small cell lung cancer (NSCLC), PA triggers ROS-mediated DNA damage, leading to cell cycle arrest and apoptosis, while overcoming drug resistance and reducing cancer stem-like properties (17). Network pharmacology and enrichment analyses further revealed that PA exerts therapeutic effects on GC primarily through the modulation of key signaling pathways, including MAPK, PI3K/AKT, and NF–κB (18). Notably, NF-κB signaling has been consistently implicated in PD–L1 upregulation and immune escape in GC [1925].

On the basis of these insights, we hypothesized that PA may modulate the immune microenvironment of GC. In this study, we investigated the effects of PA on malignant phenotypes and immune evasion mechanisms in GC cell lines. We demonstrate that PA suppresses NF-κB signaling, downregulates PD-L1 expression, and inhibits tumor immune escape. Our results provide new evidence supporting the potential of PA as a candidate agent for gastric cancer immunotherapy.

## 2 MATERIALS AND METHODS

### 2.1 Materials

SureScript^TM^ First-Strand cDNA Synthesis Kit (Cat: QP056T; GeneCopeia, USA), 2×SYBR Green qPCR Master Mix (None ROX) (Cat: G3320-05; Servicebio, China), TriQuick Reagent (Cat: R1100, Solarbio, China), MKN-45 (CL-0292, Procell, China), HGC-27 (CL-0107, Procell, China), GES-1 (CL-0563, Procell, China), Cell Counting Kit-8 (Cat: G4103, Servicebio, China), Patchouli alcohol (Cat: MUST-24053118, Purity: 99.63%, Must Bio-Technology, China), Multiskan FC Microplate Photometer (DNM-9602, Perlang, China), inverted microscope (OPTOP OD630K, Sunny Hengpin, China), chemiluminescence imaging system (ChemiScope6100, Clin, China), microplate reader (RMR-2208LZN; Ranhui, China), Invitrogen Power Blotter (Model PB0010; Thermo Fisher Scientific, China), anti-NF-kB p65 antibody (ab16502; abcam, UK), NF-κB p65 polyclonal antibody (Cat: 10745-1-AP, Proteintech, China), recombinant anti-PD-L1 antibody [EPR19759] (ab213524, Abcam, UK), PD-L1/CD274 (C-terminal) polyclonal antibody (Cat: 28076-1-AP; Proteintech, China), primary and secondary antibody diluent for immunostaining (Cat: 36206ES76; YEASEN, China), goat anti-rabbit IgG H&L (Alexa Fluor^®^ 488) (ab150077; abcam, UK), fluorescence microscope (BH2-RFCA; OLYMPUS, Japan), Annexin V-FITC/PI apoptosis kit (Cat: AP101-100-kit, MultiSciences, China), BD FACSCalibur^TM^ Flow Cytometer (E97501093, BD Biosciences, USA), Human Tumor Necrosis Factor Alpha (TNF-α) ELISA Kit (JL10208, Jianglai Biological, China), Human Interferon (IFN-γ) detection kit (JL12152, Jianglai Biological, China), TUNEL Apoptosis Detection Kit-FITC (50T) (MK1027, BOSTER, China), Dual Luciferase Reporter Gene Assay Kit (Cat: RG009, Beyotime, China), Plasmid Mini Preparation Kit (BioDev-Tech, China), pmirGLO (Honeke, China), restriction endonuclease (Thermo, USA), and GloMax^®^ 96 Microplate Luminometer (Promega, USA).

### 2.2 Methods Cell culture

Human gastric mucosal epithelial cells (GES-1), gastric cancer cell lines (AGS, BGC823, NUGC-4, SGC7901, MKN-45, HGC-27), and MFC cells were purchased from Procell Systems. The cells were maintained in DMEM supplemented with 10% FBS and 1% penicillin‒streptomycin. The cell lines were cultured at 37 °C in a humidified 5% CO_2_ incubator.

### CCK-8 assay

Gastric cancer cells in the logarithmic growth phase were seeded into 96-well plates at a density of 2000 cells per well (100 μl per well). Six replicate wells were established for each group. After the cells were cultured under 5% CO_2_ at 37 °C for 12 hours, they were divided into six different groups with increasing concentrations (0, 0.1, 0.25, 0.5, 1, 10, 20, and 50 mM), and a blank group and a solvent control group were also established. After the cells were incubated for 24, 48 or 72 hours, the culture medium was discarded. Following the instructions, cell viability was determined using a CCK-8 kit.

### Western blotting assay

Proteins were extracted from gastric cancer cells and tumor tissues using RIPA lysis buffer. The protein samples were separated by SDS‒PAGE and subsequently transferred onto PVDF membranes. Blocking was performed with 5% BSA for 2 hours. Following three washes with TBST, the membranes were incubated with specific primary antibodies overnight at 4 °C. The following day, the membranes were incubated with secondary antibodies. After three additional TBST washes, protein detection was carried out using an ECL system. Finally, ImageJ software was used to analyze the gray values and determine the relative protein expression levels.

### RT‒qPCR analysis

Total RNA was extracted from gastric cancer cells or tissues using TRIzol and subsequently reverse-transcribed into cDNA utilizing RT Master Mix, following the manufacturer’s protocol. Quantitative real-time PCR (qRT‒PCR) analysis was performed using an ABI 7500 detection system with SYBR Green (Takara) as the fluorescent dye. GAPDH served as the internal reference control gene, and the relative expression levels of the target genes were quantified using the 2^−ΔΔCt^ method (26). The primer sequences are listed in Table 1.

**Table 1.**
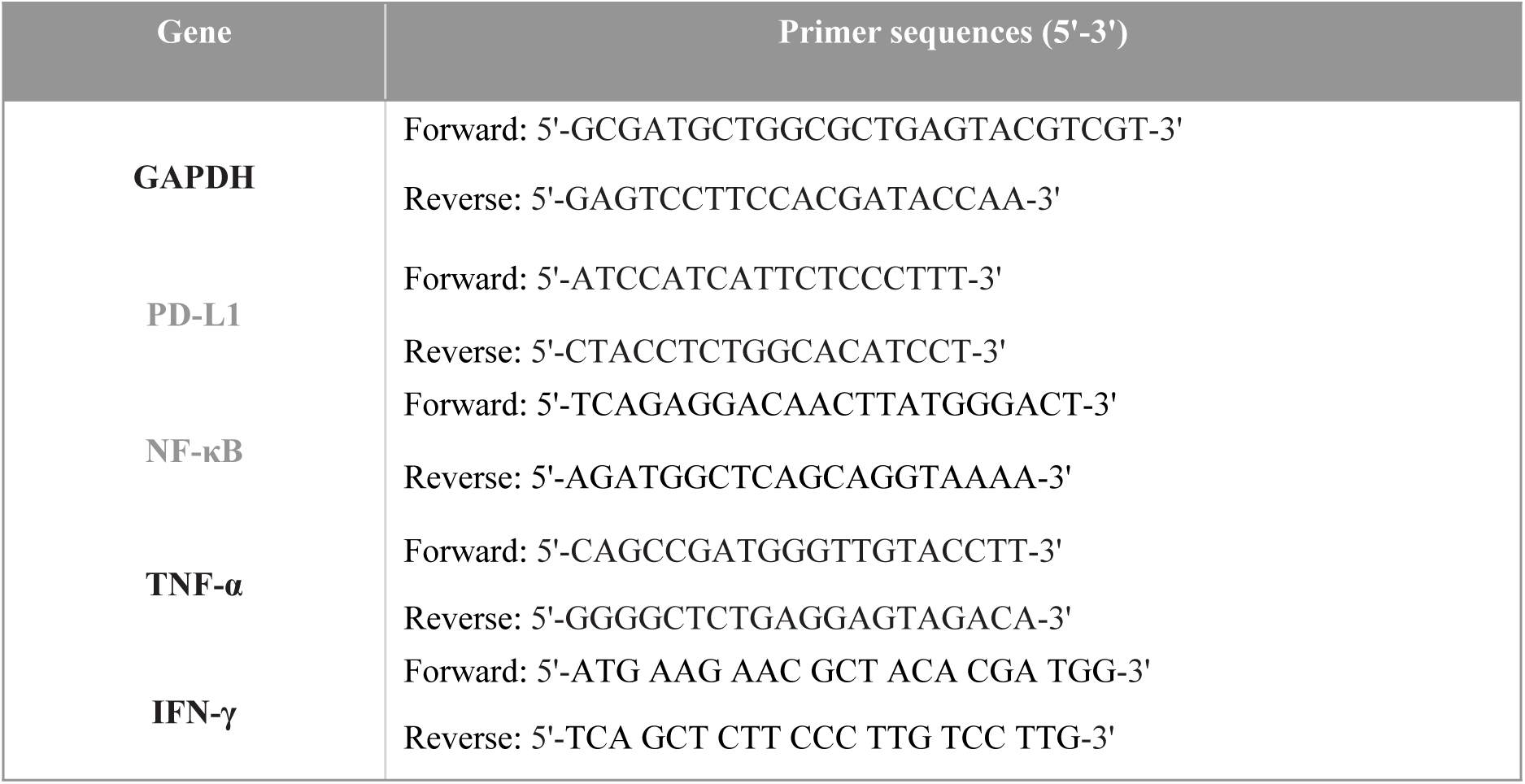
Primer sequences.

### ELISA kit

In accordance with the kit instructions, the concentrations of interferon-gamma (IFN-γ), tumor necrosis factor-alpha (TNF-α), lactate dehydrogenase (LDH) and granzyme in the different groups were detected using an ELISA kit.

### Flow cytometry

To analyze tumor-infiltrating lymphocytes (TILs), single-cell suspensions were prepared from transplanted tumor tissues by mechanical dissociation followed by enzymatic digestion with 1 mg/mL collagenase IV for 60 minutes at 37 °C. The resulting cell suspension was filtered through a 70-μm cell strainer and subjected to Percoll density gradient centrifugation (1000×g, 20 min) to isolate viable lymphocyte fractions. Following adjustment of the cell density to 1×10⁶ cells/mL in phosphate-buffered saline (PBS), 100-μL aliquots were incubated with fluorochrome-conjugated anti-mouse CD3-FITC and CD8-PE antibodies for 30 minutes at 4 °C in the dark, using appropriate isotype controls. After two washes with PBS, the cells were resuspended in 300 μL of staining buffer and analyzed using a BD FACSCanto II flow cytometer. FlowJo software was used to identify and quantify the CD3^+^CD8^+^ T-cell population.

### Apoptosis analysis

Cells from different groups were collected, washed with PBS, and resuspended in 1× binding buffer. Annexin V-FITC and PI were added, and the samples were incubated for 10–20 minutes at room temperature in the dark before being analyzed by flow cytometry. The data were processed with FlowJo software, and the cells were categorized as viable, early apoptotic, or late apoptotic/necrotic. In the Annexin V (X-axis) *vs*. PI (Y-axis) dot plot, the quadrants define cell states as follows:

Lower Left (LL: Annexin V⁻/PI⁻) = viable cells,

Lower Right (LR: Annexin V⁺/PI⁻) = early apoptotic cells,

Upper Right (UR: Annexin V⁺/PI⁺) = late apoptotic/necrotic cells,

Upper Left (UL: Annexin V⁻/PI⁺) = necrotic/mechanically damaged cells.

Total apoptosis is calculated as [(LR + UR)/Total Cells]×100%, with compensation adjusted via single-stain controls and negative boundaries set using untreated cells.

### T-cell-dependent cytotoxicity assay

Peripheral blood mononuclear cells (PBMCs) were isolated from peripheral blood and cocultured for 48 hours with gastric cancer cells (PBMCs: GC=10:1) pretreated with various concentrations (0, 0.25, 0.5, or 1 mM) of patchouli alcohol (PA) in 96-well plates. After 48 hours of cultivation, tumor cell apoptosis, cytotoxic activity, and T-cell activation were subsequently evaluated using the assays described in the following sections.

### NF-κB Overexpression and Knockdown

The NF-κB overexpression vector (pGL3-NF-κB) and NF-κB-targeting shRNA lentiviral vector (shR-NF-κB) were synthesized and constructed by BGI (Shenzhen, China). For transfection, cells were transfected with 2 μg of plasmid using Lipofectamine 3000 reagent according to the manufacturer’s protocol. After transfection for 48 h, transfection efficiency was validated by RT-qPCR.

### *In vivo* experiments

A total of 56 C57BL/6 mice (6 weeks old) were obtained from Beijing Sipeifu Biotechnology Co., Ltd. The mice were randomly assigned to seven experimental groups and housed in a specific pathogen-free environment with a 12-hour light/dark cycle. Food and water were freely available. All procedures involving animals were conducted in accordance with the guidelines approved by the Institutional Animal Care and Use Committee. MFC cells with stable NF-κB overexpression (OE-NF-κB) or stably transfected with shRNA targeting NF-κB (shRNF-κB) were established as described above. For tumor induction, 2×10^6^ MFC cells were injected subcutaneously on the dorsal side of each mouse. The experimental groups were designated as follows: control group, shR-NF-κB group, OE-NF-κB group, PA (20 mg/kg), PA (40 mg/kg), PA (20 mg/kg) + OE-NF-κB , and PA (40 mg/kg) + OE-NF-κB . Each group received the corresponding treatment, whereas the control group received an equivalent volume of normal saline. The tumor size and body weight were recorded every two days throughout the experimental period. After 14 days of treatment, the mice were euthanized via an isoflurane overdose (4% induction followed by 5% maintenance in an anesthesia chamber until the heart arrest), and the tumor tissues were excised for further analysis.

### HE staining

After the mice were euthanized, tumor tissues were collected. After being embedded in paraffin, the samples were cut into 4-μm sections and stained with hematoxylin and eosin. After the slides were sealed with neutral gum, they were observed under an optical microscope, and randomly selected fields of view were selected for imaging.

### TUNEL Assay of Frozen Sections

Slides were precoated with poly-L-lysine prior to use. Frozen tissue sections were fixed in 4% paraformaldehyde in PBS (0.01 M, pH 7.0–7.6) for 40 minutes, followed by brief washes in PBS and water, and were subsequently maintained in labeling buffer to preserve hydration. For TUNEL labeling, TdT and DIG-d-UTP were mixed in the labeling buffer and applied to the sections, which were then incubated at 37 °C for 2 hours in a humid chamber. After being washed with TBS (0.01 M), the slides were blocked and sequentially incubated with biotinylated anti-digoxigenin antibody and SABC-FITC conjugate, each of which was diluted 1:100 in SABC diluent at 37 °C, interspersed with washes using TBS.

### Immunofluorescence staining

Gastric cancer cells were seeded in 24-well plates, fixed with 1% paraformaldehyde, permeabilized with 0.1% Triton X-100, and blocked with 5% BSA. Primary antibodies against NF-κB were applied, and the samples were incubated overnight at 4 °C. On the following day, the cells were incubated with fluorescently labeled secondary antibodies for 30 minutes, followed by nuclear counterstaining with DAPI. Finally, the samples were examined and imaged under a fluorescence microscope. We used Image J software to quantify NF-κB expression in MKN-45 and HGC-27 cells.

### PD-L1 promoter luciferase reporter construction

The full-length synthesized promoter sequence was inserted into the pGL3-Basic vector via the KpnI and HindIII restriction sites to construct the pGL3-PD-L1-WT plasmid. The pGL3-PD-L1 mutant plasmid was generated from the wild-type pGL3-PD-L1 template using a site-directed mutagenesis kit according to the manufacturer’s instructions.

### Luciferase analysis

Gastric cancer cells in the logarithmic growth phase from each experimental group were seeded into 48-well plates at 20,000 cells per well, with three technical replicates per group. After 24 hours, transfection reagent was added, and the cells were transfected for 48 hours. The ratio of firefly luciferase activity to Renilla luciferase activity was calculated for each cellular sample.

### Statistical analysis

The data are presented as the mean±SD. Statistical analyses were performed using SPSS software (version 22.0). Differences among multiple groups were analyzed by one-way ANOVA followed by LSD post hoc testing. Comparisons between two groups were conducted using paired Student’s t tests. Statistical significance was defined as *P* < 0.05.

## 3 RESULTS

### 3.1 Inhibitory effects of PA on the viability of gastric cancer cells and the expression of PD-L1 and NF-κB

First, the mRNA expression levels of NF-κB and PD-L1 were detected in gastric cancer cell lines. RT‒qPCR analysis revealed significant overexpression of both NF-κB and PD-L1 mRNA in gastric cancer cells compared with the normal gastric mucosal cell line GES-1 (Fig. 1A). Among the tested lines, HGC-27 and MKN-45 cells exhibited relatively high mRNA levels and were therefore selected for further investigation. Following treatment with different concentrations of PA (0, 0.1, 0.25, 0.5, 1, 10, 20, and 50 mM), CCK-8 assays demonstrated a concentration- and time-dependent decrease in gastric cancer cell viability (Fig. 1B). Furthermore, RT‒qPCR and western blotting analyses consistently revealed that compared with the untreated control, PA treatment significantly reduced both the mRNA (Fig. 1C) and protein (Fig. 1D) expression levels of NF-κB and PD-L1. These results indicate that PA can inhibit gastric cancer cell proliferation and downregulate key molecules involved in immune escape.

**Figure 1.**
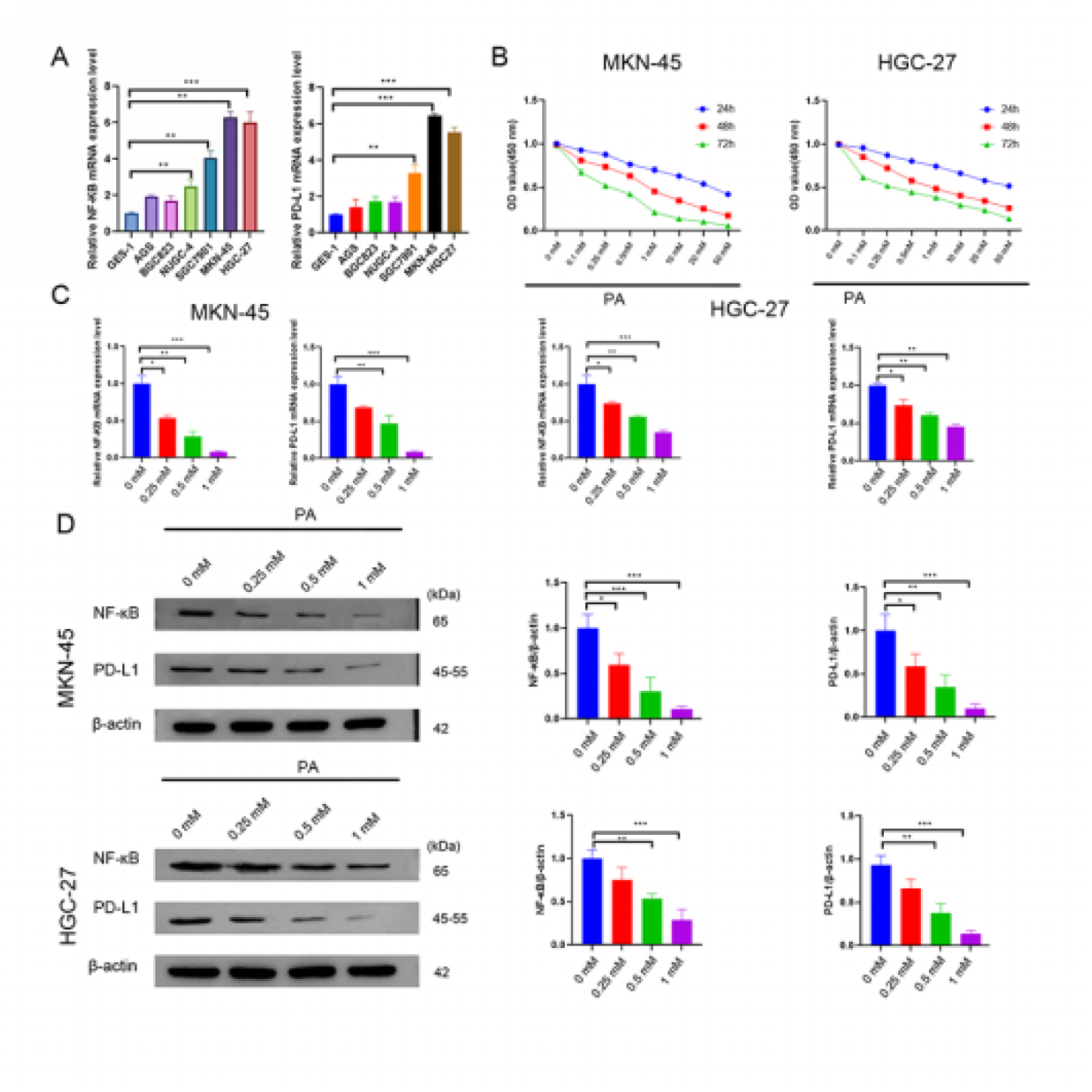
Inhibitory effects of PA on the viability of gastric cancer cells and the expression of PD-L1 and NF-κB. **(A)** mRNA expression levels of NF-κB and PD-L1 in human gastric mucosal cells and gastric cancer cell lines. **(B)** Viability of the gastric cancer cell lines HGC-27 and MKN-45 treated with increasing concentrations of PA (0 mM, 0.1 mM, 0.25 mM, 0.5 mM, 1 mM, 10 mM, 20 mM and 50 mM) for 24 hours, 48 hours, and 72 hours. **(C)** Effect of PA treatment on the mRNA expression of NF-κB and PD-L1 in HGC-27 and MKN-45 cells. **(D)** Effect of PA treatment on the protein expression of NF-κB and PD-L1 in HGC-27 and MKN-45 cells. \**P* < 0.05, \*\**P* < 0.01, *** *P*< 0.001; n=3 independent experiments.

### 3.2 PA pretreatment sensitizes gastric cancer cells to PBMC-mediated cytotoxicity through the activation of CD8⁺ T cells

To evaluate the immunomodulatory effects of PA, gastric cancer cells were pretreated with PA for 48 hours prior to coculture with PBMCs. Flow cytometry analysis demonstrated that PA pretreatment significantly enhanced the PBMC-induced apoptosis of both HGC-27 and MKN-45 cells in a dose-dependent manner (Fig. 2A). LDH release assays confirmed that compared with the control treatment, PA pretreatment markedly increased the cytotoxic activity of PBMCs against gastric cancer cells (Fig. 2B). Immunophenotyping of cocultured PBMCs by flow cytometry revealed a significant increase in the proportion of cytotoxic CD3⁺CD8⁺ T cells following PA pretreatment (Fig. 2C). Notably, the frequency of IFN-γ-producing CD8⁺ T cells (IFN-γ⁺CD8⁺) also increased in a PA dose-dependent manner (Fig. 2D), indicating enhanced functional activation. This immunostimulatory profile was further supported by cytokine marker analysis. The results of the ELISA analysis revealed that compared with the control group, the coculture supernatants from the groups pretreated with PA contained significantly elevated levels of IFN-γ, TNF-α, and granzyme B. Taken together, these findings indicate that PA pretreatment sensitizes gastric cancer cells to immune attack by promoting a CD8⁺ T-cell-activating microenvironment, thereby enhancing PBMC-mediated tumor cell death.

**Figure 2.**
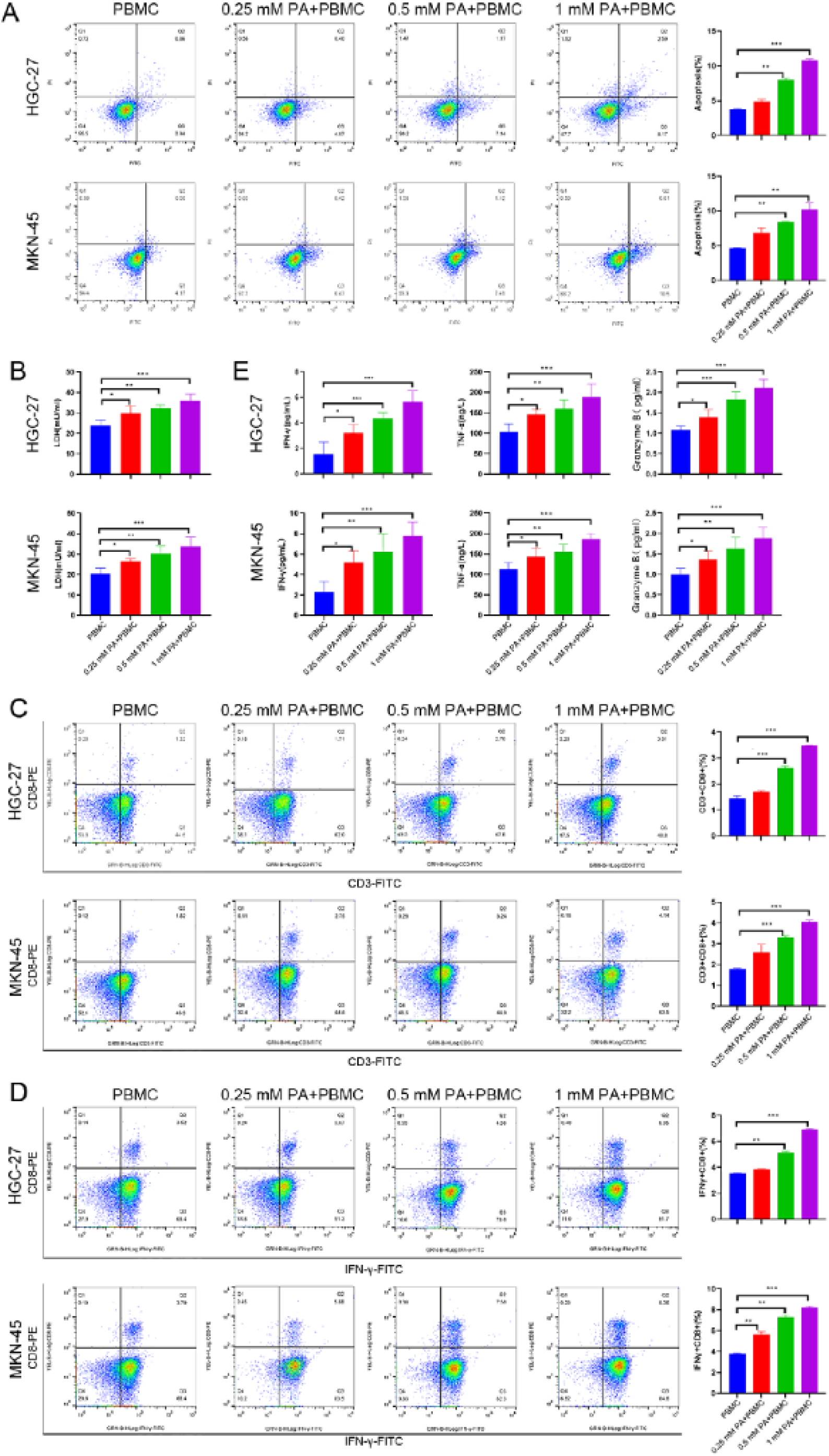
PA pretreatment sensitizes gastric cancer cells to PBMC-mediated death through the activation of CD8⁺ T cells. **(A)** Apoptosis of HGC-27 and MKN-45 cells analyzed by flow cytometry. **(B)** Analysis of the cytotoxic activity of PBMCs against gastric cancer cells by LDH release assays and concentrations of IFN-γ, TNF-α, and granzyme B in the coculture supernatant measured by ELISA. **(C, D)** Immunophenotypic analysis of the cocultured PBMCs. Flow cytometry was used to quantify the proportions of CD3⁺CD8⁺ T cells (C) and IFN-γ–producing CD8⁺ (IFN-γ⁺CD8⁺) T cells **(D)**. \**P* < 0.05, \*\**P* < 0.01, \*\*\**P* < 0.001, *vs.* the control group; n=3 independent experiments.

### 3.3 PA suppresses NF-κB-induced PD-L1 upregulation in gastric cancer cells

Western blotting analysis revealed that, compared with those in the control group, the protein levels of AKT, NF-κB, and PD-L1 in the group treated with either an shRNA targeting NF-κB (shR-NF-κB) or 0.5 mM PA were significantly lower. The inhibitory effects of 0.5 mM PA and the shR-NF-κB on NF-κB and PD-L1 expression were similar; in both MKN-45 and HGC-27 cells, the NF-κB overexpression (OE-NF-κB) group markedly upregulated AKT, NF-κB, and PD-L1 expression. Notably, in MKN-45 cells cotreated with 0.5 mM PA and the OE-NF-κB group, the expression levels of AKT, NF-κB, and PD-L1 were significantly lower than those in the OE-NF-κB-alone group. In HGC-27 cells, cotreatment with 0.5 mM PA group also significantly reduced the expression levels of NF-κB and PD-L1 compared with those in the OE-NF-κB-alone group (Fig. 3A), indicating that PA can counteract the protein induction triggered by NF-κB activation. The RT‒qPCR results demonstrated a consistent decreasing trend in NF-κB mRNA expression (Fig. 3B). Immunofluorescence staining further confirmed a marked reduction in NF-κB nuclear localization in both the PA-treated and shR-NF-κB groups compared with that in the control group, whereas NF-κB nuclear translocation was significantly enhanced in the OE-NF-κB group. Furthermore, compared with OE-NF-κB group alone, PA + OE-NF-κB cotreatment markedly suppressed NF-κB nuclear translocation (Fig. 3C, D). Immunofluorescence staining revealed that PA inhibited the nuclear translocation of NF-κB and suppressed its activation of downstream signaling molecules. Collectively, these findings suggest that PA suppresses NF-κB-driven PD-L1 upregulation in gastric cancer cells.

**Figure 3.**
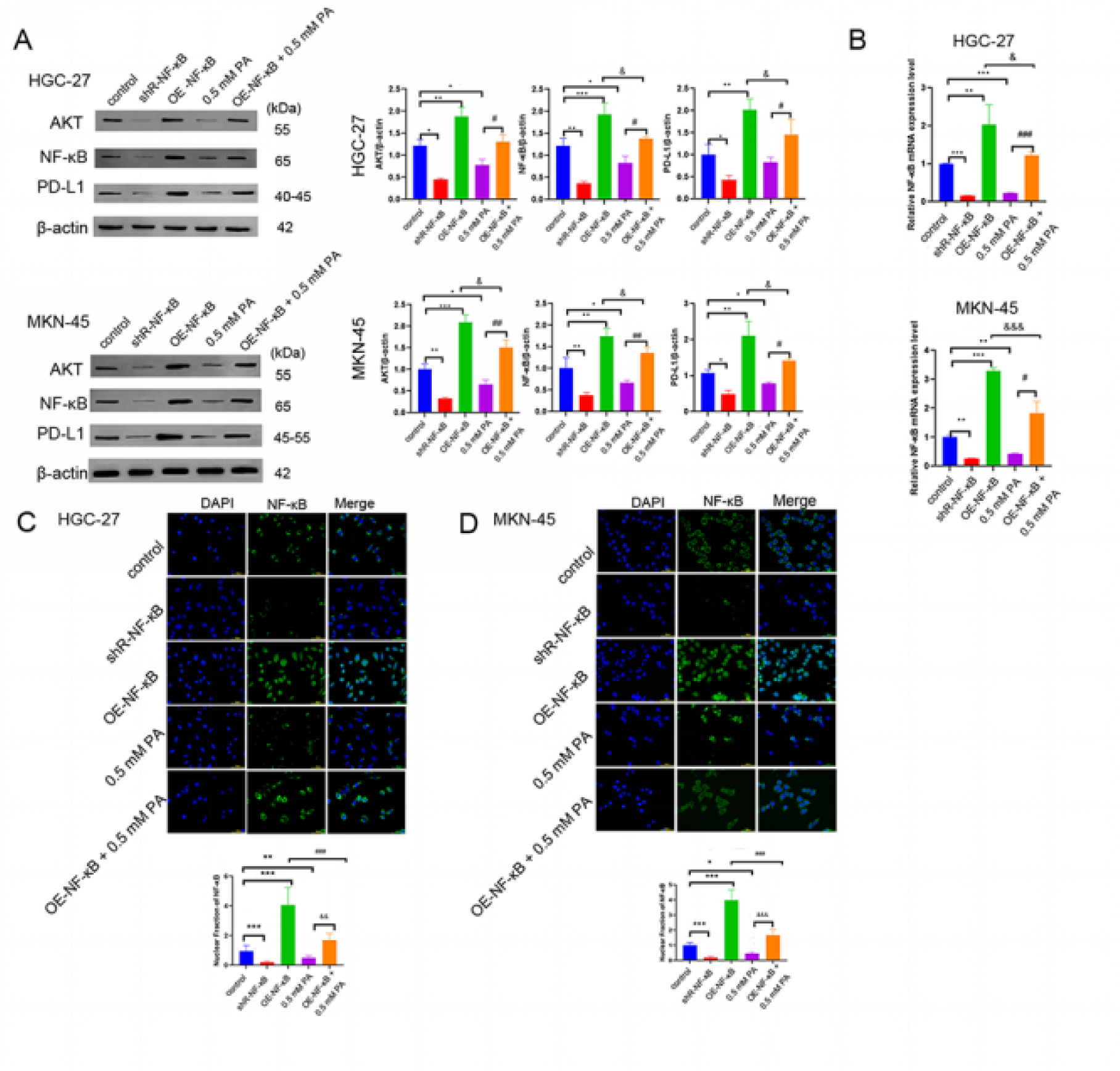
Patchouli alcohol suppresses NF-κB-induced PD-L1 upregulation in gastric cancer cells. **(A)** western blotting analysis of AKT, NF-κB, and PD-L1 protein expression in HGC-27 and MKN-45 cells treated as indicated. **(B)** RT‒qPCR analysis of NF-κB mRNA expression in HGC-27 and MKN-45 cells under various treatments. **(C, D)** Immunofluorescence staining was used to measure the levels of NF-κB in HGC-27 **(C)** and MKN-45 **(D)** cells; OE-NF-κB: overexpression of NF-κB; shR-NF-κB: shRNA targeting NF-κB. Scale bar, 100 μm. \**P* < 0.05, \*\**P* < 0.01, \*\*\**P* < 0.001, *vs.* the control group; ^#^*P* < 0.05, ^##^*P* < 0.01, ^###^*P* < 0.001, *vs.* the OE-NF-κB+0.5 mM PA group; ^&^*P* < 0.05, ^&&&^*P* < 0.001, *vs.* the OE-NF-κB group; n=3 independent experiments.

### 3.4 PA inhibits tumor growth and induces apoptosis in a mouse model of gastric cancer

To evaluate the *in vivo* antitumor effect of PA, C57BL/6 mice were subcutaneously injected with MFC cells with stable NF-κB overexpression or stably transfected with shRNA targeting NF-κB to establish a transplanted tumor model. After 14 days of treatment, the tumors were harvested for analysis. As shown in Fig. 4A-C, compared with the control treatment, both the shR-NF-κB group and the PA treatment significantly suppressed transplanted tumor growth, as evidenced by the reduced tumor size (Fig. 4A), tumor volume (Fig. 4B), and final tumor weight (Fig. 4C). Histopathological examination by HE staining revealed that tumors from PA-treated mice exhibited reduced cellularity, disorganized architecture, and extensive necrotic areas (Fig. 4D). In contrast, coadministration of an OE-NF-κB group with PA partially reversed these morphological changes, resulting in tumors with higher cellular density and more preserved structure than those in the PA monotherapy groups (Fig. 4D). Notably, the necrotic areas in the combination groups remained more prominent than those in the control group, with the high-dose PA (40 mg/kg) plus OE-NF-κB group showing more necrosis than the low-dose (20 mg/kg) combination group did (Fig. 4D). Consistent with the observed tissue damage, the TUNEL assay demonstrated a significant increase in apoptotic cells within the tumor tissues of both the shR-NF-κB group and the PA treatment group (Fig. 4E). This pro-apoptotic effect of PA was attenuated when it was combined with the OE-NF-κB, as the combination groups showed lower levels of apoptosis than their respective PA-alone groups did (Fig. 4E). Taken together, these *in vivo* results indicate that PA inhibits tumor growth, induces apoptosis, and causes significant intratumoral damage, effects that are mediated, at least in part, through the suppression of NF-κB signaling. No obvious tissue damage or abnormalities were observed in these organs, indicating that 40 mg/kg PA did not cause apparent toxicity.

**Figure 4.**
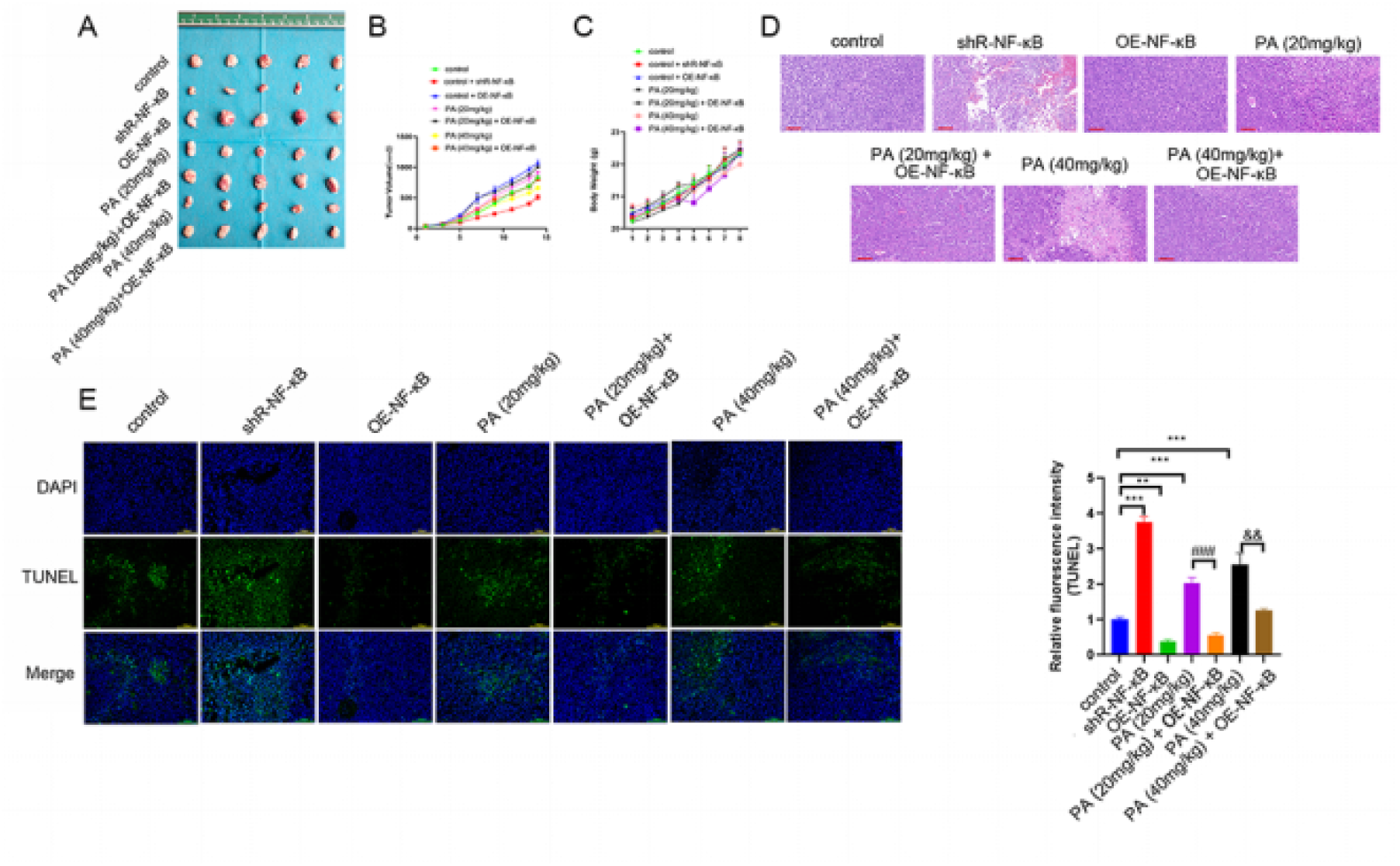
PA inhibits tumor growth and induces apoptosis in a mouse model of gastric cancer. **(A)** Representative images of tumors excised from each treatment group at the endpoint. **(B)** Dynamic changes in tumor volume during the 14-day treatment period. **(C)** The final tumor weight was measured at the time of sacrifice. **(D)** Representative HE-stained sections of tumor tissues. Scale bar, 100 μm. **(E)** Detection of apoptotic cells in tumor tissues by TUNEL staining. Green fluorescence indicates TUNEL-positive (apoptotic) cells. Nuclei are counterstained with DAPI (blue). OE-NF-κB: overexpression of NF-κB; shR-NF-κB: shRNA targeting NF-κB. Scale bar, 100 μm. The data in B, C, and E are presented as the mean ± SD (n=5 mice per group). \*\**P* < 0.01, \*\*\**P* <001, *vs.* the control group; ^##^*P* < 0.01, *vs.* the PA (20 mg/kg) group; ^&&^*P* < 0.01, *vs.* the PA (40 mg/kg) group.

### 3.5 PA inhibits the NF-κB/PD-L1 axis and reverses immune evasion in a mouse model of gastric cancer

To evaluate the *in vivo* effect of PA on the NF-κB/PD-L1 axis, the tumor tissues described in Section 3.4 were analyzed. Immunofluorescence staining revealed a significant reduction in NF-κB expression in tumors from the shR-NF-κB and PA treatment groups compared with the control group, whereas NF-κB expression was markedly increased in the OE-NF-κB group (Fig. 5A). Western blotting analysis yielded consistent results, with significantly lower NF-κB protein levels in the shR-NF-κB and PA groups than in the control group (Fig. 5B). To assess the functional consequences of modulating this axis by PA, we analyzed the tumor immune microenvironment by flow cytometry. Compared with those in the control group, the proportions of both tumor-infiltrating CD3⁺CD8⁺ T cells and IFN-γ⁺CD8⁺ T cells were significantly greater in the shR-NF-κB and PA (40 mg/kg) treatment groups (Fig. 5C, D). Moreover, compared with the OE-NF-κB group, the PA (40 mg/kg) + OE-NF-κB cotreatment group exhibited significantly greater proportions of CD3⁺CD8⁺ T cells and IFN-γ⁺CD8⁺ T cells (Fig. 5C, D). Collectively, these results demonstrate that PA inhibits the NF-κB/PD-L1 signaling axis *in vivo*, thereby reversing tumor-mediated immune evasion in a mouse model of gastric cancer.

**Figure 5.**
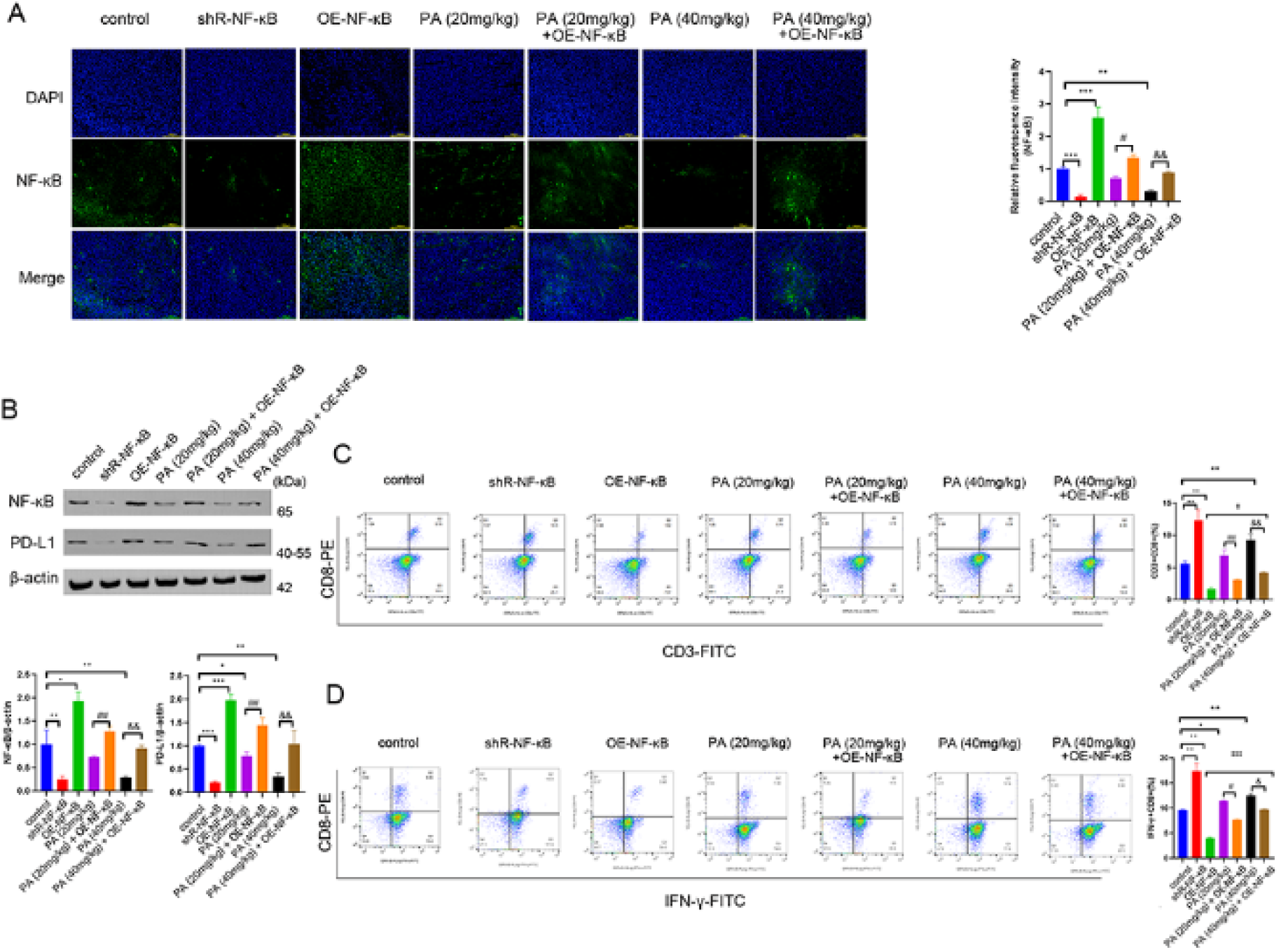
PA inhibits the NF-κB/PD-L1 axis and reverses immune evasion in a mouse gastric cancer model. Mice bearing MFC allografts were treated as indicated for 14 days. **(A)** Representative immunofluorescence images showing NF-κB expression (green) in tumor tissues. Nuclei are counterstained with DAPI (blue). Scale bar, 100 µm. **(B)** The protein expression of NF-κB and PD-L1 in GC cells was evaluated using western blotting. **(C, D)** The proportions of CD3^+^CD8^+^ **(C)** and IFN-γ^+^CD8^+^ **(D)** T cells were determined by flow cytometry. OE-NF-κB: overexpression of NF-κB; shR-NF-κB: shRNA targeting NF-κB. \**P* < 0.05, \*\**P* < 0.01, \*\*\**P* < 0.001, *vs*. the control group; ^#^*P* < 0.05, ^##^*P* < 0.01, *vs*. the PA (20 mg/kg) group; ^&^ *P* < 0.05, ^&&^*P* < 0.01, *vs*. the PA (40 mg/kg) group; ^‡^*P* < 0.05, ^‡‡‡^*P* < 0.001, *vs*. the OE-NF-κB group; n=3 independent experiments.

### 3.6 PA inhibits PD-L1 transcription via the suppression of NF-κB activity

Building on the evidence that PA downregulates NF-κB and PD-L1 expression, we next investigated whether NF-κB directly regulates PD-L1 transcription. Luciferase reporter assays were performed using a construct containing the wild-type PD-L1 promoter (PD-L1-WT) in gastric cancer cells after validation of efficient NF-κB overexpression or knockdown (Fig. 6A, B). As shown in Fig. 6C, NF-κB overexpression significantly increased luciferase activity driven by the PD-L1-WT promoter in both HGC-27 and MKN-45 cells, whereas knockdown of NF-κB substantially reduced luciferase activity. To confirm the specificity of NF-κB binding, we employed a promoter construct with a mutated NF-κB binding site (PD-L1-MUT). As demonstrated in Fig. 6D, mutation of this site completely abrogated the responsiveness of the PD-L1 promoter to either NF-κB overexpression or knockdown. These results demonstrated that NF-κB directly binds to the PD-L1 promoter to activate its transcription. Taken together, our data indicate that PA downregulates PD-L1 expression through direct suppression of NF-κB–mediated transcriptional activation.

**Figure 6.**
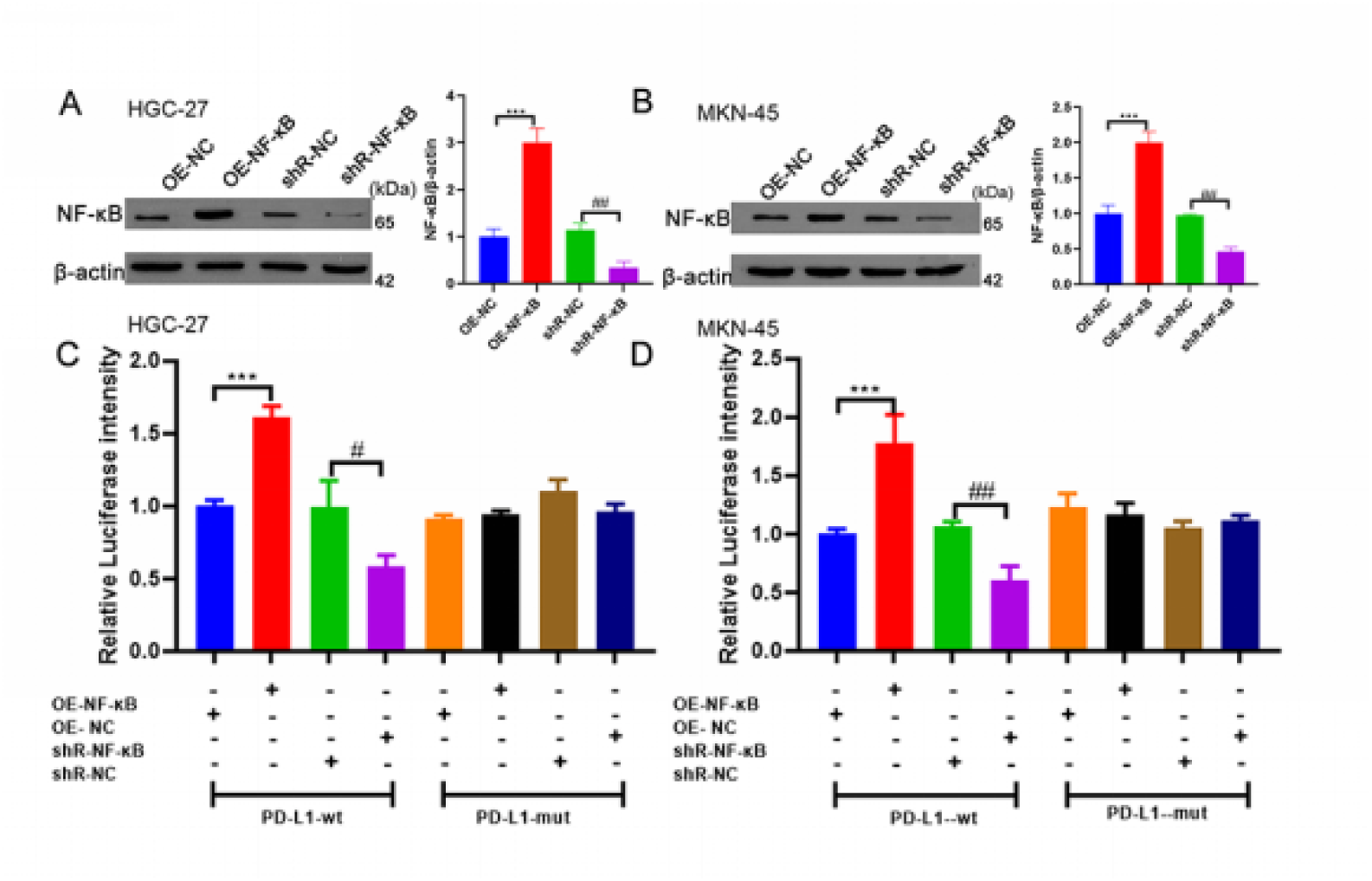
NF-κB directly regulates PD-L1 promoter activity in gastric cancer cells. **(A, B)** Validation of NF-κB overexpression or knockdown in MKN-45 cells and HGC-27 cells. **(C)** Luciferase reporter assays in HGC-27 cells. **(D)** Luciferase reporter assays in MKN-45 cells. OE-NF-κB: overexpression of NF-κB; OE-NC: overexpression negative control; shR-NF-κB: shRNA targeting NF-κB; shR-NC: shRNA negative control. \*\*\**P* < 0.001, \**P* < 0.05, *vs.* the OE-NC group; ^#^*P* < 0.05, ^##^*P* < 0.01, *vs.* the shR-NC group; n=3 independent experiments.

## 4 DISCUSSION

Despite significant advances in therapeutic strategies, gastric cancer remains a major global public health challenge because of its high mortality rate. Accumulating evidence (27, 28) indicates that immune evasion plays a pivotal role in gastric cancer progression and significantly compromises the efficacy of conventional treatments such as chemotherapy and radiotherapy. Therefore, targeting immune evasion mechanisms may represent a crucial approach for improving gastric cancer therapy. Among the key mediators of immune evasion, the PD-1/PD-L1 axis has been extensively documented in gastric cancer (29, 30). In this study, we provide *in vitro* and *in vivo* evidence that PA suppresses gastric cancer progression by inhibiting the NF-κB pathway and downregulating PD-L1 expression.

Our previous work using network pharmacology and molecular docking revealed that PA, a bioactive component of Pogostemon cablin, exerts anti-gastric cancer effects via the NF-κB pathway. Given that NF-κB is a well-established regulator of PD-L1 overexpression in tumor cells, we sought to determine whether PA could modulate PD-L1 expression through NF-κB inhibition. We confirmed that PA inhibits NF-κB nuclear translocation, thereby reducing PD-L1 expression, with an inhibitory potency that is comparable to that observed after shRNA-mediated NF-κB silencing.. *In vitro* cytotoxicity assays demonstrated that PA enhances T-cell–mediated tumor cell death, thereby mitigating immune evasion. Importantly, even under conditions of NF-κB overactivation, PA induces apoptosis and necrosis in gastric cancer cells in a dose-dependent manner. Specifically, compared with the medium-dose group, the high-dose PA + OE-NF-κB group exhibited more extensive necrosis, indicating that PA maintains dose-dependent cytotoxic effects despite heightened NF-κB signaling. This detailed elucidation of the PA-mediated PD-L1 downregulation constitutes a novel contribution of our study.

It is important to recognize that NF-κB regulates tumor immunity beyond the PD-1/PD-L1 axis. Growing evidence implicates it in modulating other immune checkpoints, such as TIM-3 and CD155. TIM-3 impairs the function of various immune cells and represents a promising immunotherapy target (31). In hepatocellular carcinoma, TIM-3 can activate NF-κB to promote immune evasion (32), and conversely, TNF-α upregulates TIM-3 on NK cells via NF-κB (33), highlighting a bidirectional regulatory interplay. Similarly, CD155 (PVR), a ligand for TIGIT and CD226, is a “next-generation” checkpoint widely expressed in solid tumors. Its coexpression with PD-L1 is linked to poor outcomes ([3437]). NF-κB hyperactivation enhances CD155 transcription and stability, promoting the activity of the TIGIT-CD155 immunosuppressive axis (38, 39). Although direct evidence in gastric cancer is limited, the pronounced inflammatory nature of this disease strongly suggests that NF-κB serves as a critical link between chronic inflammation and an immunosuppressive microenvironment, potentially by upregulating the expression of molecules such as CD155 and TIM-3. This conceptual framework guides our future research into whether PA can remodel the immune landscape by targeting these additional pathways.

The multitarget mechanism of PA aligns with the traditional Chinese medicine (TCM) principle of “strengthening healthy qi and eliminating pathogenic factors”. Its “strengthening” effect is evidenced by the increased CD3⁺CD8⁺ T-cell proportions and increased IFN-γ secretion, akin to the immune-boosting properties of TCM agents such as *Astragalus membranaceus* (40, 41). Conversely, its “pathogen-eliminating” effect occurs through direct induction of cancer apoptosis and potent inhibition of the protumorigenic NF-κB inflammatory pathway. This dual action corresponds to the TCM strategy of “clearing heat and detoxifying,” which aims to restore homeostasis by suppressing pathological inflammation. Thus, the integrated approach of PA, which involves simultaneously modulating immune escape and directly killing tumor cells, exemplifies the distinct advantages of multitarget TCM therapies.

## 5 CONCLUSION

Our study demonstrated that PA exerts antitumor effects on gastric cancer by inhibiting NF-κB signaling, downregulating PD-L1 expression, and enhancing T-cell cytotoxicity (summarized in Fig. 7). These findings not only provide a mechanistic basis for the application of PA but also highlight its potential as a multitarget agent bridging direct antitumor activity and immune microenvironment modulation.

**Fig 7.**
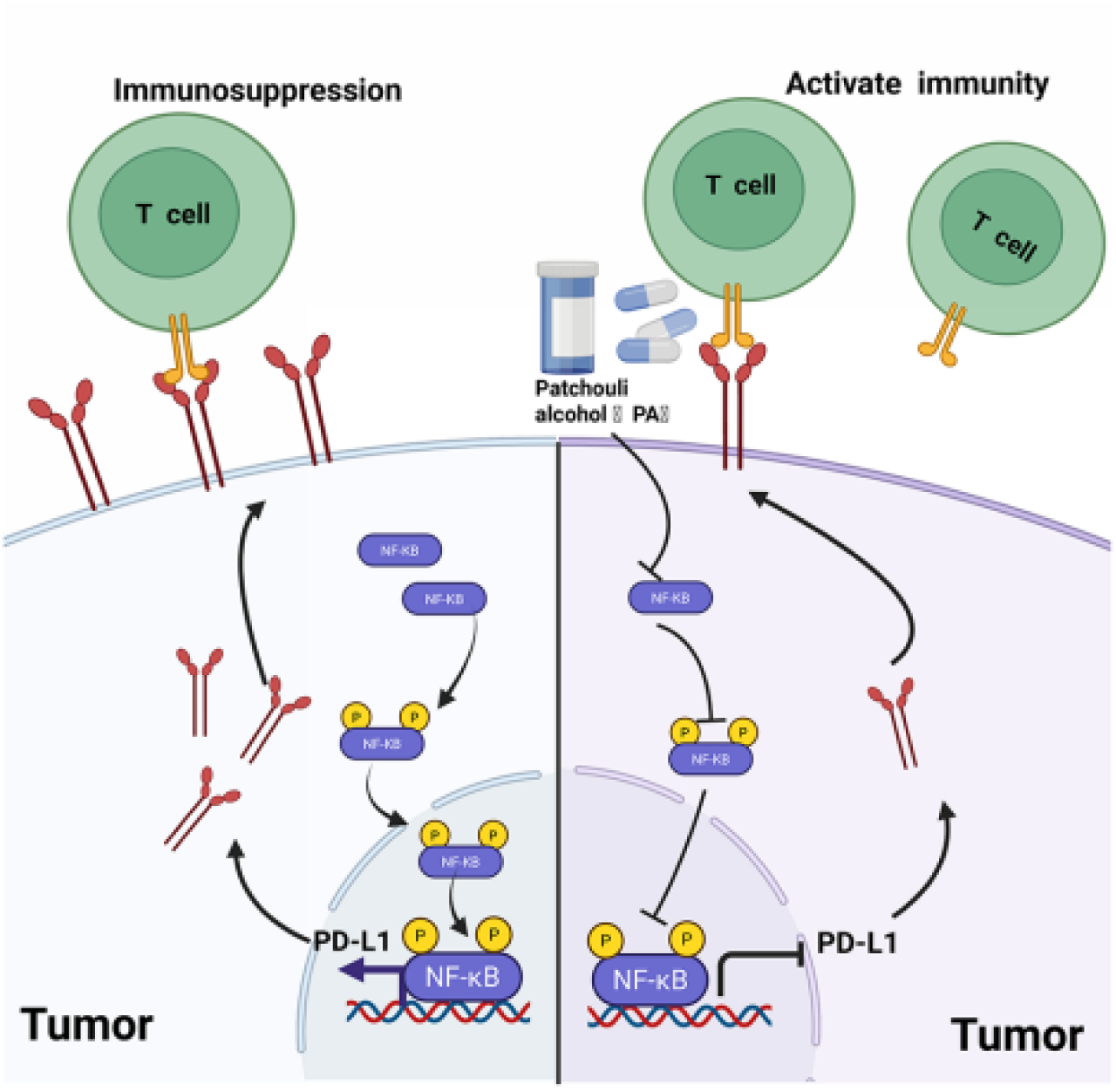
In gastric cancer cells, after NF-κB is activated, it can translocate into the nucleus and directly bind to the promoter region of the PD-L1 gene, promoting its transcription and thereby upregulating the expression of PD-L1, which facilitates immune escape by gastric cancer cells. PA can inhibit the expression and activation of NF-κB, weakening its positive regulatory effect on PD-L1 transcription and significantly reducing the protein level of PD-L1, thereby alleviating the T-cell function inhibition mediated by PD-L1 and enhancing the specific killing effect of CD8⁺ T cells on gastric cancer cells.

## DATA AVAILABILITY STATEMENT

The original contributions presented in the study are included in the article/Supplementary Material; further inquiries can be directed to the corresponding author.

## AUTHOR CONTRIBUTIONS

HK and WZF conceived and designed the study. HK, HQ and YH collected and analyzed the data and performed the statistical analysis. HK wrote the article. HK, HQ and YH prepared the pictures and tables. HK, WZF, FL and LHW provided suggestions and participated in the revision of the article. All the authors read and approved the final manuscript.

## FUNDING

This study was financially supported by the Natural Science Foundation of Inner Mongolia (2023QN08003), the Project of Inner Mongolia Medical University (YKD2022LH004), the High-Level Clinical Specialty Development Technology Project for Public Hospitals of the Inner Mongolia Autonomous Region (2023pp005), Green Seed Talent Program of Peking University Cancer Hospital Inner Mongolia Hospital (QM202308), the Science and Technology Project of the Public Hospital Research Joint Fund of Inner Mongolia Medical Sciences Academy (2024GLLH0386, 2025GLLH0208), the National Natural Science Foundation of China (82074144, 82460865), the Inner Mongolia Autonomous Region Science and Technology Planning Project (2022YFHH0095, 2023KJHZ0023), the Scientific and Technological Innovative Research Project for Inner Mongolia Universities (NMGIRT2327), the Natural Science Foundation of Inner Mongolia (2024ZD31), the Capital’s Funds for Health Improvement and Research (2022-2-1024), the Project of Inner Mongolia Medical University (YKD2023TD010, YKD2024TD014, YKD2023QN010), the High-Level Clinical Specialty Development Technology Project for Public Hospitals in Inner Mongolia Autonomous Region (2023SGGZ072, 2023SGGZ074 (07), 2023SGGZ114, 2023pp001), and the Green Seed Talent Program of Peking University Cancer Hospital Inner Mongolia Hospital (QM202309).

## CONFLICT OF INTEREST

The authors declare that the research was conducted in the absence of any commercial or financial relationships that could be construed as potential conflicts of interest.

## Notes

### Competing Interest Statement

The authors have declared no competing interest.

